# OAC-PCA: orthogonal adjustment of confounding effects in principal component analysis for metabolomics data mining

**DOI:** 10.64898/2026.05.21.726783

**Authors:** Misaki Kurata, Hiroyuki Yamamoto, Hiroshi Tsugawa

**Affiliations:** Department of Biotechnology and Life Science, Tokyo University of Agriculture and Technology, 2-24-16 Naka-cho, Koganei, Tokyo, 184-8588 Japan; Human Metabolome Technologies, Inc., 246-2 Mizukami, Kakuganji, Tsuruoka, Yamagata 997-0052, Japan; Graduate School of Pharmaceutical Sciences, Kyoto University, 46-29 Yoshida-Shimoadachi-cho, Sakyo-ku, Kyoto 606-8501, Japan

## Abstract

Principal component analysis (PCA) is widely used in mass spectrometry-based metabolomics for exploratory data mining. Statistical testing of loading values can extract metabolite features associated with score patterns, but this approach requires principal components (PCs) to remain orthogonal while loadings are defined as correlation coefficients between PC scores and variables. Adjustment for Confounding PCA (AC-PCA) was previously developed to explore biologically meaningful components from data matrices affected by biological and technical confounders. However, AC-PCA does not simultaneously ensure PC orthogonality and a correlation-coefficient definition of loadings, limiting the statistical interpretation of its loadings. Here, we reformulated AC-PCA as Orthogonal Adjustment for Confounding effects in PCA (OAC-PCA). In OAC-PCA, PCs remain orthogonal, and loadings retain this correlation-coefficient interpretation. These properties enable statistical testing of metabolite associations while accounting for confounding effects.

## Introduction

In the analysis of multidimensional omics data, feature extraction by dimensionality reduction methods, such as principal component analysis (PCA) and partial least squares (PLS), is widely used^1,2^. PCA identifies linear combinations of variables that maximize variance in a reduced-dimensional space^3^. In metabolomics, PCA score plots are commonly used to visualize high-dimensional data in two- or three-dimensional subspaces. Variables associated with these score patterns are then interpreted using loadings along the principal component axes. Because principal components (PCs) are orthogonal, loading values can be subjected to statistical hypothesis testing, enabling variables to be selected based on statistical criteria rather than subjective inspection. This property helps reduce the risk of misleading biological interpretation^4,5^.

However, omics data are often affected by confounding factors derived from technical variation, such as batch effects, and biological variation unrelated to the primary research question^6,7^. Batch effects refer to technical variation unrelated to the biological objective. They can arise from differences in sample processing protocols, experimental conditions and data acquisition platforms^8^. Such effects are widely observed across omics fields, including genomics, RNA-seq, single-cell RNA-seq, proteomics, metabolomics, and multi-omics studies. Biological confounders, in contrast, arise from intrinsic characteristics such as age, sex, and ethnicity and are particularly relevant in cohort studies^9^. Both technical and biological confounders can obscure genuine biological signals and lead to misinterpretation.

To address this issue, Adjustment for Confounding PCA (AC-PCA) was proposed to extract biologically meaningful components from data matrices affected by confounders^10^. AC-PCA has two formulations. The first is a general formulation that can be solved as an eigenvalue problem, but its loadings cannot be expressed as correlation coefficients. Therefore, their statistical properties remain unclear. The second formulation incorporates an L1 constraint to improve the interpretability of PCs. Even when this problem is simplified by considering only an L2 constraint for confounders, it requires solving a generalized eigenvalue problem, which does not guarantee the orthogonality of the loading vectors. Thus, the statistical properties of the loadings in this formulation also remain unclear.

To overcome these limitations, we reformulated AC-PCA as Orthogonal Adjustment for Confounding effects in PCA (OAC-PCA). OAC-PCA introduces two auxiliary variables, t and s: s is penalized for confounder-related variation, whereas t is defined analogously to the standard PCA score vector. This formulation allows the weight vectors to be obtained by solving an eigenvalue problem, thereby preserving their orthogonality. In addition, the loadings can be interpreted as correlation coefficients between the PC scores of the auxiliary variable s and each metabolite feature, making them statistically interpretable. We evaluated OAC-PCA using one simulated dataset and two real datasets and compared its performance with PCA and AC-PCA.

## Results

### Case 1: Application to simulated data

We first compared PCA, AC-PCA, and OAC-PCA using simulated data from the AC-PCA R package (https://github.com/linzx06/AC-PCA). This dataset was designed to mimic confounded brain data, consisting of three donors, 10 brain locations, and 400 variables (see the detail in **Materials and Methods**). PCA produced score distributions dominated by donor-specific differences, obscuring the biological pattern associated with the 10 brain locations. In contrast, both AC-PCA and OAC-PCA recovered this latent biological pattern after adjusting for donor effects (**Figure 1**).

**Figure 1.**
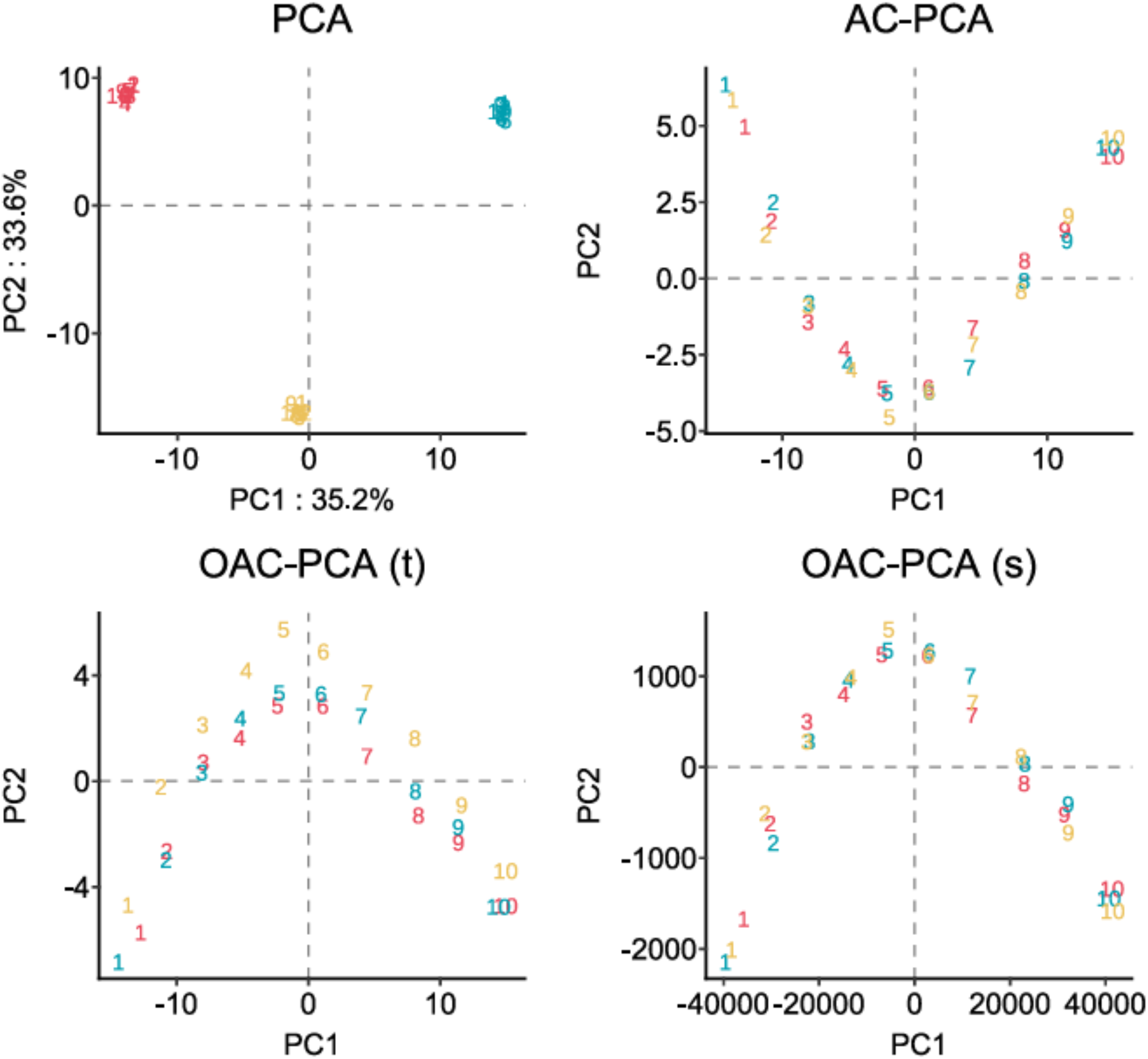
Comparison of PC score distributions. Scatter plots of the first and second PC scores calculated by PCA, AC-PCA, and OAC-PCA. For OAC-PCA, scores are shown for the auxiliary variables *t* and *s*. Colors represent the three donors, and numbers 1–10 indicate the simulated brain locations.

### Case 2: Application to single-cell RNA sequencing data

We evaluated OAC-PCA by applying it to mice single-cell RNA sequence data^11^. This dataset contains CD45^⁺^ cells isolated from lung tumor tissues of lung cancer mice using the KP1.9 tumor model. It was originally generated to investigate the role of tumor-infiltrating myeloid cells (TIMs) in tumor progression. The dataset includes three batches and seven major immune cell clusters: T cells, natural killer (NK) cells, dendritic cells, B cells, monocytes, macrophages, dendritic cells, basophils, and neutrophils. These clusters were annotated using a Bayesian cell classifier^12,13^ based on marker gene information. The public dataset had been filtered to remove low-quality cells, including cells with high mitochondrial gene expression, followed by doublet removal and gene expression normalization. We further preprocessed the dataset using the standard workflow implemented in the Seurat R package. For this study, we selected two batches, round1_20151128 and round2_20151217, and two cell clusters, T cells and B cells. The round1_20151128 batch contained 579 T cells and 641 B cells, whereas the round2_20151217 batch contained 434 T cells and 879 B cells. The final dataset consisted of 2,533 cells and 10,000 genes.

We then performed PCA and OAC-PCA (**Figure 2A**). PCA separated the two cell clusters but also captured variation attributable to batch effects. In contrast, OAC-PCA reduced the influence of batch effects while preserving the separation between T cells and B cells. In the OAC-PCA score plot, T cells were distributed toward the positive direction of PC1 and the negative direction of PC2, whereas B cells showed the opposite pattern. We interpreted the OAC-PCA results using loading values. Statistical hypothesis testing of the loadings extracted genes that significantly contributed to PC1 and PC2 (**Figure 2B**). After false discovery rate correction using the Benjamini–Hochberg method^14^, 2,268 genes were significant for PC1 and 1,945 genes were significant for PC2, with q-value < 0.05. Genes contributing to the negative direction of PC1 and the positive direction of PC2 included T cell– associated genes such as Cd28, a receptor widely expressed on T cells during antigen presentation^15^; Cd4, a co-receptor involved in T cell activation and a well-known marker of helper T cells; and Cd8, a co-receptor that binds the T cell receptor (TCR) and MHC class I molecules and serves as a marker of cytotoxic T cells^16^. Other top-ranked genes included Thy-1, the first identified T cell marker^17^, along with many T cell–specific genes. These results indicate that this axis reflects T cell identity and function. These results indicate that these axes are characterized by genes predominantly associated with T cell identity and function. Conversely, genes contributing to the positive direction of PC1 and the negative direction of PC2 included major B cell marker genes such as Cd19 and Cd20 (Ms4a1), which are involved in B cell differentiation and maturation^18^, and Cd79a, which encodes a component of the B cell receptor complex^19^. Genes encoding MHC class II molecules, including Cd74^20^, H2-Eb1, and H2-Aa^21^, were also ranked among the top contributors. Because MHC class II molecules are expressed by antigen-presenting cells, including macrophages, dendritic cells, and B cells^22^, this axis likely reflects B cell identity and antigen presentation. These results are consistent with known differences between T cell and B cell immune systems.

**Figure 2.**
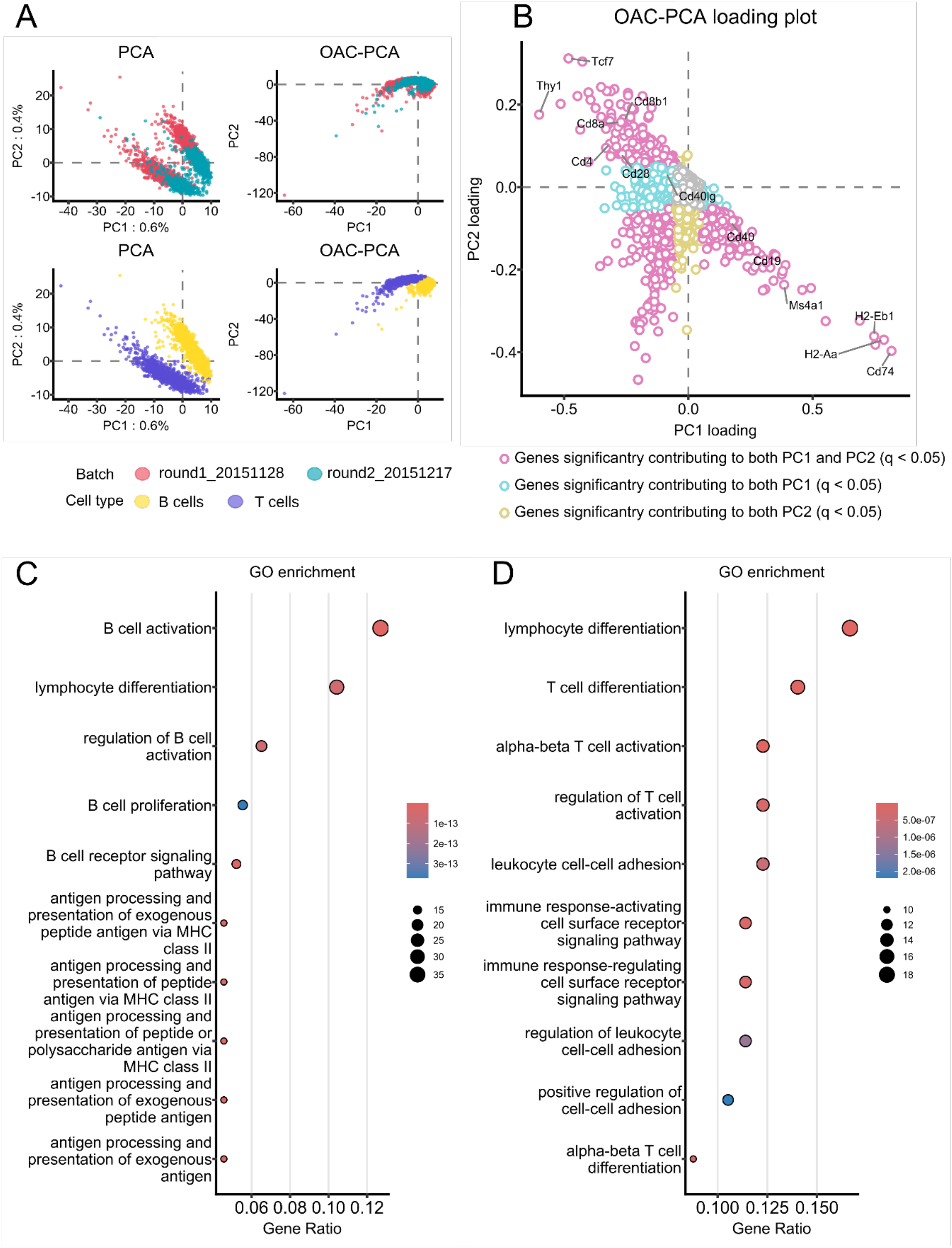
Evaluation of OAC-PCA using single-cell RNA sequencing data. (A) Scatter plots of PC1 and PC2 scores obtained by PCA (left) and OAC-PCA (right). OAC-PCA scores are represented by the auxiliary variable *t*. The upper panels are colored by batch (round1_20151128 and round2_20151217), whereas the lower panels are colored by cell cluster (T cells and B cells). (B) Distribution of loading values for 10,000 genes. Loading values for PC1 and PC2 are plotted, with genes significantly contributing to both PC1 and PC2, PC1 only, or PC2 only highlighted in color (q-value < 0.05). (C) Gene Ontology (GO) enrichment analysis for PC1. The plot shows the top ten significantly enriched terms among genes that contributed positively to PC1. (D) GO enrichment analysis for PC2. The plot shows the top ten significantly enriched terms among genes that contributed positively to PC2.

We next performed enrichment analysis to characterize the gene sets contributing to each principal component. Genes with q-value < 0.05 and positive contributions to PC1 were subjected to Gene Ontology (GO) analysis. The same procedure was applied to genes contributing positively to PC2. Among the top ten ontology terms enriched for genes positively contributing to PC1, we observed B cell–related terms, including “B cell activation,” “regulation of B cell activation,” “B cell proliferation,” and MHC class II–related terms such as “antigen processing and presentation of exogenous peptide antigen via MHC class II” (**Figure 2C**). These results indicate that PC1 characterizes B cell signatures, particularly antigen-presenting capacity. In contrast, genes positively contributing to PC2 were enriched for T cell–related terms, including “T cell differentiation,” “alpha-beta T cell activation,” and “regulation of T cell activation” (**Figure 2D**). Thus, PC2 captures functional signatures of T cells. Taken together, these results demonstrate that OAC-PCA mapped the distinct biological signatures of B cells and T cells into PC1 and PC2, respectively.

### Case 3: Application to age-related lipidomics data

We next applied OAC-PCA to an age-related lipidomics data from mice kidneys (9 weeks, 12 months, 18 months, and 24 months) ^23^. At each time point, samples were obtained from both sexes (male/female) under two microbial conditions, specific-pathogen-free (SPF) and germ-free (GF), with four biological replicates per condition. Conventional PCA was dominated by sex differences, and age-associated signals were not clearly resolved. In contrast, OAC-PCA accounted for sex as a confounder and captured age-dependent lipidomic changes (**Figure 3A**). PC1 separated 9-week-old mice from aged mice, suggesting that this axis reflects lipidome alterations associated with postnatal maturation. PC2 showed a positive association with age, indicating progressive age-related changes (**Figure 3B**). We then calculated loading values and performed statistical testing to identify lipid species contributing to PC1 and PC2 (**Figure 3C**). Among 527 detected lipid species, 84 lipids showed significant positive correlations with PC1, whereas 163 showed significant negative correlations.

**Figure 3.**
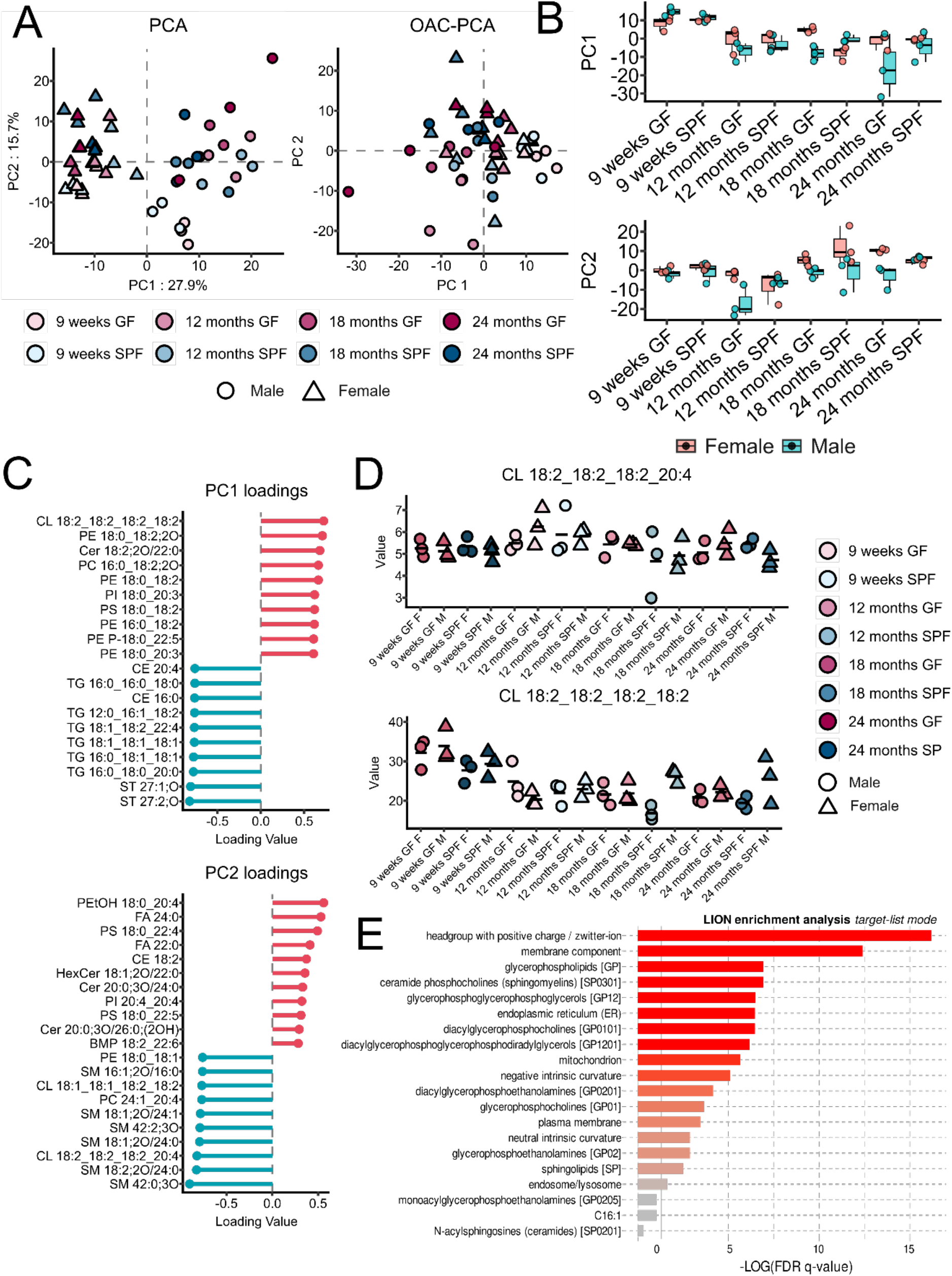
Evaluation of OAC-PCA using age-related lipidomics data from mouse kidneys. (A) Scatter plots of PC1 and PC2 scores obtained by PCA (left) and OAC-PCA (right). OAC-PCA scores are represented by the auxiliary variable *t*. Circles and triangles indicate male and female mice, respectively. Samples are colored by microbial status: red for germ-free (GF) mice and blue for specific-pathogen-free (SPF) mice. Age groups (9 weeks, 12 months, 18 months, and 24 months) are indicated by color intensity, with darker shades representing older ages. (B) Box plots of PC1 (upper) and PC2 (lower) scores. Points are colored by sex. PC1 captures the lipidomic profile characteristic of the 9-week-old stage, whereas PC2 reflects progressive age-associated changes. (C) Top 10 positive and negative loading values for PC1 and PC2. Lollipop charts show lipid species with the largest positive and negative loadings along each PC axis. (D) Lipid ontology enrichment analysis. Lipids showing significant negative correlations with PC2 (q-value < 0.05) were analyzed using the LION/web target mode. All detected lipid species were used as the background list. (E) Age-related changes in the detected abundances of tetralinoleoyl-CL (CL 18:2_18:2_18:2_18:2) and CL 18:2_18:2_18:2_20:4. Red and blue points represent GF and SPF mice, respectively, with color intensity indicating age.

Notably, tetralinoleoyl-cardiolipin (CL 18:2_18:2_18:2_18:2) showed the highest loading score for PC1. Cardiolipin is a structurally unique phospholipid with a dimeric architecture, four acyl chains, and two negative charges. It is mainly localized in the mitochondrial inner membrane, where it supports the structural organization and functional activity of proteins involved in mitochondrial energy metabolism^24^. CL 18:2_18:2_18:2_18:2, which contains four linoleic acid chains, is the most abundant cardiolipin species in the mammalian heart. This linoleoyl-rich composition is critical for maintaining cytochrome c oxidase activity (respiratory chain complex IV) and mitochondrial respiratory capacity^25^. The abundance profile of tetralinoleoyl-CL matched the PC1 score pattern, peaking at 9 weeks, declining by 12 months (**Figure 3D**). Conversely, lipids negatively correlated with PC1, which were enriched in aged mice, were mainly storage lipids, including triacylglycerols (TG), sterols (ST), and cholesterol esters (CE). These findings suggest that the 9-week stage represents a lipidome state associated with active mitochondrial oxidative metabolism, which gradually shifts toward lipid storage and metabolic stagnation with age.

For PC2, six lipids showed significant positive correlations and 142 showed significant negative correlations (q-value < 0.05). The positively correlated lipids included HexCer 18:1;2O/22:0, a long-chain hexosylceramide reported to accumulate in the kidney during aging.^26^ For the 142 lipids negatively correlated with PC2, enrichment analysis using LION/web^27^ highlighted terms related to the endoplasmic reticulum (ER) and mitochondria, particularly cardiolipin species (**Figure 3E**).

Notably, these cardiolipins showed a unique profile distinct from that of CL 18:2_18:2_18:2_18:2, the primary contributor to PC1: they peaked at 12 months and then decreased toward 24 months (**Figure 3D**). Age-related oxidative stress promotes the production of reactive oxygen species (ROS). Previous studies have suggested that ROS-induced cardiolipin oxidation is associated with the upregulation of *Alcat1*, an acyl-CoA–dependent acyltransferase that catalyzes cardiolipin remodeling using monolysocardiolipin (MLCL) and dilysocardiolipin as substrates at mitochondria-associated membranes (MAMs). This remodeling can promote ROS-sensitive cardiolipin species and contribute to mitochondrial dysfunction^28^. Thus, the distinct cardiolipin profiles observed in this study may reflect the lipidome alterations associated with mitochondrial remodeling in aging. In summary, OAC-PCA revealed mitochondria-associated lipid metabolism in kidney where mitochondrial inner membrane lipid properties begin to change as early as 12 months of age.

## Materials and Methods

### Simulated data

This dataset mimics the brain data. 3 donors and 400 variables and 10 locations, and the value was generated as below.

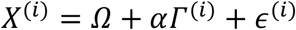

Ω represents the low-rank shared component across donors, *Γ*^(*i*)^ captures donor-specific variation, and *ϵ*^(*i*)^denotes Gaussian noise. The goal of this simulation is to capture the shared low-rank pattern Ω across donors.

### Data preprocessing

The single cell RNA-seq data was preprocessed based on the standard workflow provided by the Seurat R package. Highly variable genes were identified, and the top 10,000 most variable genes were selected. In this study, two batches (round1 _20151128 and round2 _20151217) and two cell clusters (T cells and B cells) were selected. In round1_20151128, the dataset contained 579 T cells and 641 B cells, while in round2_20151217, it contained 434 T cells and 879 B cells, resulting in a total of 2533 cells. The lipidomics data used to evaluate OAC-PCA was processed as the follow’s methods. The original dataset contained 884 lipid species. To reduce noise influence, lipids were filtered based on the following criteria: signal-to-noise ratio (S/N) average > 10, mean intensity greater than or equal to twice the blank value, and coefficient of variation (CV) ≤ 2. In addition, lipid species with odd-numbered fatty acyl chains were excluded. After filtering, 708 lipids remained. When duplicate lipid species were present, the entry with the higher average intensity was retained, resulting in a final dataset of 527 unique lipid species.

#### AC-PCA

AC-PCA in general form is formulated as:

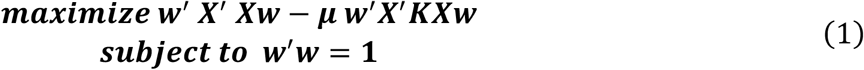

where X is a mean-centered *N* × *p* data matrix (*N*: sample size, *p*: variable size like metabolites or genes); *K* = *Y* ′*Y. Y* is a mean-centered *N* × *l* confounder matrix (*l*: the number of confounders), so *w*′*X*′*KXw* represent the confounders variation; w is a vector of weights

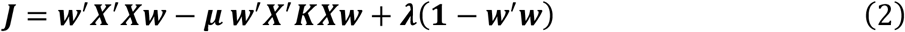

where λ is Lagrange multiplier. By differentiating *J* partially with *w*, it can be expressed as the following equation.

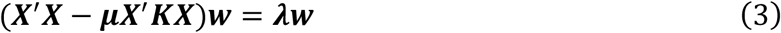

Then, *w* can be determined as eigen problem. However, in this formulation, the loadings can be expressed as follows, but their direct relationship with *w* remains unclear.

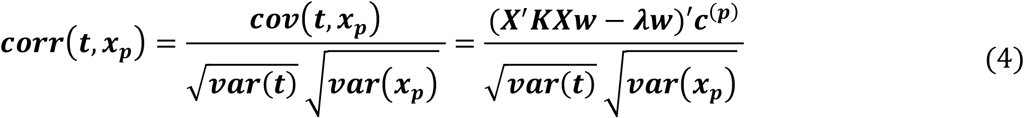

The alternative version of AC-PCA adding L2 constraint is formulated as:

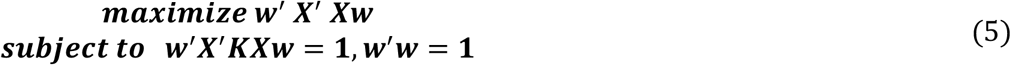

Letting λ be the Lagrange multiplier, the Lagrangian function can be expressed as:

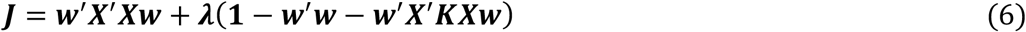

By differentiating *J* partially with *w*, it can be expressed as the following equation.

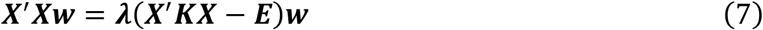

Here, *E* denotes the identity matrix. In this case, ***w*** can be determined by solving a generalized eigenvalue problem. This formulation does not preserve orthogonality among PCs, which hampers statistical testing and interpretation of loadings like general form.

#### OAC-PCA

OAC-PCA is formulated as:

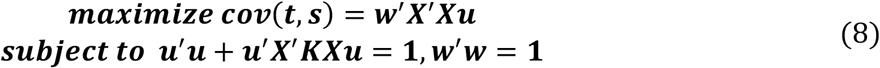

This formulation is similar to that of OS-PCA^29^.

Using the Lagrange multiplier method, we reformulated it as follows.

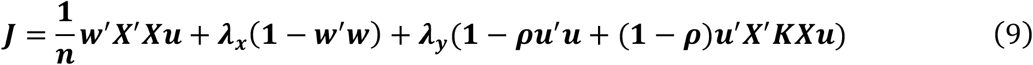

Where *λ*_*x*_ and *λ*_*y*_ are Lagrange multipliers and *ρ* is tuning parameter controls the strength of regularization. When *ρ* is 0, this formulation coincides with classical PCA. By differentiating *J* partially with respect to *w* and *u*, respectively, and transforming, it can be expressed as the following two equations:

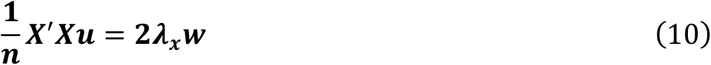

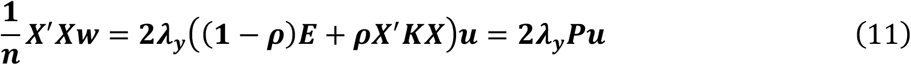

Then, these can be rewritten as the following two eigenvalue problems.

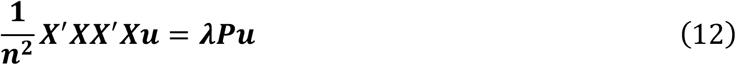

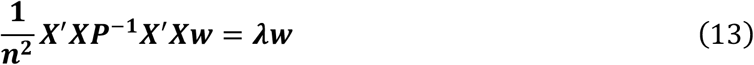

where *λ* = 4*λ*_*x*_*λ*_*y*_.

### Statistical property of OAC-PCA loading for auto-scaled data

In the interpretation of PCA results, value of the loading is important. In PCA, loading is generally defined as the correlation coefficients between variables and eigenvectors. Consequently, by applying it to statistical hypothesis testing, it is possible to select variables that are statistically significant. In OAC-PCA, loading was defined as follows.

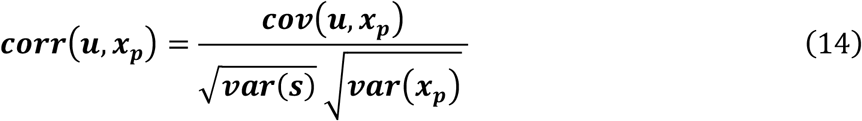

When X is scaled data, *var*(*x*_*p*_) = 1.

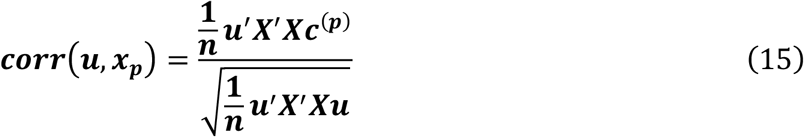

From Equation (10), 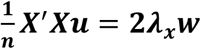, it can be expressed as below.

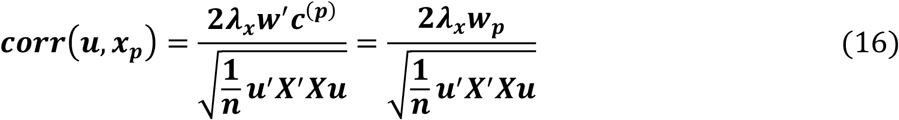

The diameter is not affected by p-th variables. So, the correlation coefficient between *u* and *xp* correlates with *w* by correlation coefficient 1. Using the loading *r* defined, the following t-statistic can be calculated, which follows a t-distribution with n-2 degrees of freedom^4^.

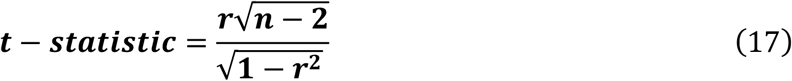

By applying a t-test, the *p*-value can be calculated which enables variable selection based on the statistic.

## Discussion

Confounding factors can distort dimensionality reduction results and complicate biological interpretation. AC-PCA was developed to address this issue by extracting components after adjusting for confounder effects. However, AC-PCA does not simultaneously provide PC orthogonality and a correlation-coefficient definition of loadings, limiting the statistical interpretation of loading values. In this study, we reformulated AC-PCA as OAC-PCA, or Orthogonal Adjustment for Confounding effects in PCA. In OAC-PCA, PC orthogonality is preserved, and loadings can be interpreted as correlation coefficients between PC scores and variables. These properties enable hypothesis testing of variable associations while accounting for confounding effects.

Using the simulated data, we confirmed that OAC-PCA adjusts for confounder effects as effectively as AC-PCA. We then applied OAC-PCA to single-cell RNA sequencing data and age-related lipidomics data. In the single-cell RNA sequencing data analysis, OAC-PCA revealed distinct biological signatures of T and B cells that were partially obscured by batch effects in conventional PCA. In the lipidomics analysis, OAC-PCA identified age-associated lipidomic shifts in mouse kidneys, highlighting cardiolipins as major contributors. Notably, different cardiolipin acyl-chain compositions showed distinct time profiles during aging. These patterns may reflect age-associated mitochondrial remodeling, including increased ROS production and Alcat1-mediated cardiolipin lipid remodeling. Together, these findings suggest that age-related lipidomic changes in the kidney are associated with mitochondrial membrane lipid metabolism.

Overall, OAC-PCA provides a framework for revealing biological variation masked by technical and biological confounders. By selecting significant variables based on statistical criteria, this method supports downstream enrichment analysis and molecular interpretation. As demonstrated here using transcriptomics and lipidomics datasets, OAC-PCA is broadly applicable to omics data affected by confounding variation. It should be particularly useful for cohort studies with complex biological covariates and for multi-omics datasets affected by technical heterogeneity.

## Data availability

Simulated data was downloaded from AC-PCA R package (https://github.com/linzx06/AC-PCA). Mouse lung single-cell RNA-seq data were obtained from the R/Bioconductor scRNAseq package. Details are available at https://rdrr.io/github/drisso/scRNAseq/man/ZilionisLungData.html. The mice aging lipidome data were obtained from Supplementary Data 1 of the previously published article (https://www.nature.com/articles/s43587-024-00610-6).

## Code availability

OAC-PCA source code is available at https://github.com/systemsomicslab/oac-pca

## Acknowledgments

This research was supported by the Japan Science and Technology Agency (JST) ERATO (JPMJER2101 to H.T.), JST FOREST (JPMJFR230H to H.T.), JST NBDC (JPMJND2305 to H.T.), JST ASPIRE (JPMJAP2505 to H.T.), and the JSPS KAKENHI (24K02011, 24H00043, 24H00392, 24K21269, 25H01425, and 25H01426 to H.T.).

## Author Contributions

H.Y. and H.T. designed the study. H.Y. provided the idea of mathematical methodology. M.K. implemented the methodology and performed the data analysis. H.T. provided the idea of mathematics and biological insights. M.K., H.Y., and H.T. wrote the manuscript. All authors have thoroughly discussed this project and helped improve the manuscript.

